# Parallel polygenic urban adaptation despite high gene flow in a coastal marine invertebrate

**DOI:** 10.1101/2025.10.03.680100

**Authors:** Madison L. Armstrong, Katie L. Erickson, Rebecca Hawthorne, Brenda Cameron, Rachael A. Bay

## Abstract

Urbanization results in novel environments, offering a unique opportunity to investigate natural selection on small spatiotemporal scales. Although each city is different, geographically distinct cities can in some ways represent evolutionary replicates, allowing for the investigation of parallel and predictable evolutionary responses to urban environments. Additionally, there are few examples of urban adaptation in high gene flow marine systems, whose patterns might differ markedly from aquatic and terrestrial systems. Using whole genomes from Pacific purple sea urchins (*Strongylocentrotus purpuratus)* across three coastal cities spanning >2000km, we investigated genomic signals for adaptation to urban environments. We found genetic variants differentiating urban and nonurban sites within each city region, despite high gene flow and little evidence for differentiation across latitudinal gradients. While these SNP-level candidates for selection were largely non-overlapping, polygenic approaches uncover a distinct parallel signal of urban adaptation across the sampled range. Our results suggest that adaptation over small scale urbanization gradients is possible even in high gene flow systems and the polygenic architecture of adaptation is, at least in part, parallel. More broadly, our work highlights the importance of polygenic methods in ecological genomics in expanding our understanding of how evolutionary forces operate in natural systems.

## Introduction

Recently, there has been growing interest in understanding how urban environments alter the ecology and evolution of the organisms that inhabit them (Carlen et al., 2026). Urbanization fragments habitat, increases temperature, and introduces novel physical substrates and chemical stressors (i.e., pollutants) (Rivkin et al., 2019). Organisms living in urban environments can be subject to novel conditions not experienced previously in their evolutionary history. Due to this, urbanization offers an unprecedented opportunity to investigate rapid contemporary evolutionary processes in real time. Simultaneously, urban environments offer replicate testbeds for such adaptation (Diamond & Martin, 2021; Lambert et al., 2021; Santangelo, Ness, et al., 2022). However, each city is not necessarily identical, as they can each have their own uniquely heterogeneous landscape shaped by geography, culture, religion and politics of the area (Carlen et al., 2025; Dant et al., 2025; Santangelo, Roux, et al., 2022)

So far, the evidence of adaptation to urban environments has been dominated by examples in terrestrial systems (Campbell-Staton et al., 2020; Daron et al., 2026; Diamond et al., 2018; McLean et al., 2005; Murray Stoker & Johnson, 2024; Santangelo, Ness, et al., 2022; Yilmaz et al., 2021) and to a lesser extent aquatic systems (Brans et al., 2021; Kern & Langerhans, 2018). Beginning with the seminal studies of industrial melanism in peppered moths (Kettlewell, 1955), examples of genetic adaptation to urban environmental variation have been documented in a range of terrestrial and aquatic systems to increased temperature (Campbell-Staton et al., 2020; Diamond et al., 2018; McLean et al., 2005; Yilmaz et al., 2021), pollution/pesticides (Brans et al., 2021; Daron et al., 2026) and impervious surfaces (Murray Stoker & Johnson, 2024; Santangelo, Ness, et al., 2022). Few examples have been shown in marine systems (Fusco et al., 2021; Todd et al., 2019), with one dominant exception being the observation of pollution resistance in marine fish populations (Park et al., 2025; Whitehead et al., 2017). Although coastal marine environments are subjected to a many stressors resulting from urbanization, we are just beginning to understand how those environmental changes alter the ecology and evolution of marine populations.

The marine intertidal zone lies at the transition from sea to land, and thus faces a complex mixture of urban stressors, including exposure to runoff, wastewater, and coastline development (Alter et al., 2021). Wastewater pollution can result in introduction of novel synthetic chemicals in the marine environment, and these plumes have also been shown to result in decreased pH, increased water temperature, and decreased salinity (Gierach et al., 2017; Lin et al., 2025; Washburn et al., 1992). Urban wastewater can also limit larval dispersal, effectively reducing connectivity between populations (Puritz & Toonen, 2011). Alternatively, urbanization can also result in high admixture between populations due to human-mediated movement of individuals, as seen in several species of blue mussels inhabiting boat launches and ports (Simon et al., 2020). While there is a rich literature on how abiotic stressors like salinity, temperature and pH impact marine species (Gleason & Burton, 2015; González Durán et al., 2018; Griffiths et al., 2021; Hoegh-Guldberg et al., 2007; Hughes et al., 2017; Kültz, 2015; Przeslawski et al., 2015; Wong & Hofmann, 2020), less work has been done to investigate the overall impact of urbanization on marine organisms despite the clear interaction of these stressors.

Marine species often have larger ranges and experience higher gene flow than their terrestrial counterparts (Hedgecock, 1986; Hellberg, 2009; Marko & Hart, 2017), potentially limiting adaptation over the small spatial scales at which urbanization gradients are realized. However, this “myth of marine uniformity” has been a recent point of contention (Overgaard Therkildsen, 2026). A growing body of literature in marine evolution suggests that balanced polymorphisms in marine systems can lead to adaptation even over small spatial scales (Rumberger et al., 2025; Sanford & Kelly, 2011; Tepolt et al., 2022). Microgeographic adaptation, defined as local adaptation that occurs below the scale of dispersal, can occur when there is strong and repeated differential selection over small spatial scales (Levene, 1953; Richardson et al., 2014). Theory and simulations highlight that even with gene flow, if adaptation is polygenic we might expect different signals of selection among populations to the same environmental conditions (Yeaman, 2015). Marine organisms with long-distance planktonic dispersal offer an ideal study system to investigate microgeographic adaptation and urban environments offer replicate environmental gradients for such studies.

To investigate genomic signals of selection associated with urbanization, we conducted whole genome sequencing in Pacific purple sea urchins (*Strongylocentrotus purpuratus*). *S. purpuratus* is a widespread marine invertebrate species that is broadly distributed across the northeastern Pacific coastline of North America. They are found in the low intertidal and subtidal zones, ranging from Alaska to Baja California, Mexico, inhabiting a large range of rocky intertidal and kelp forest sites in both urbanized and nonurban coastal waters. *S. purpuratus* is a critically important species in California kelp forest ecosystems as a dominant consumer of kelp. Further, they are a valuable model species in genomics and ecotoxicology due to the vast genomic resources and well-recorded development stages (Armstrong, 2025; Armstrong et al., 2025; Cunningham et al., 2020; Pespeni et al., 2013; Petak et al., 2023; Rumberger et al., 2025; Sea Urchin Genome Sequencing Consortium et al., 2006). Previous studies have found no population structure from Oregon to Southern California (Pespeni et al., 2012, 2013; Pespeni & Palumbi, 2013; Rumberger et al., 2025), yet there is some evidence of adaptation to small scale environmental variation, including tidal zone (Rumberger et al., 2025) and pH (Pespeni et al., 2012, 2013; Pespeni & Palumbi, 2013).

While previous studies examined adaptation to long-standing environmental gradients, we focus on adaptation to urbanization, a much more recent selection pressure. Further, we use urban environments as a model to understand patterns of parallel adaptation in this highly dispersive marine system, implementing novel polygenic methods. To do this, we sampled from both urban and nonurban locations across three coastal urban and nearby nonurban regions spanning over 2000 km to address two questions. First, within each region, do signals of urban adaptation exist over these small spatial scales? Second, if there is evidence of urban adaptation, are those signals parallel across geographically disparate urban environments?

## Methods

### Site Classification

A paired urban/nonurban site design was implemented across three Eastern Pacific coastal cities: Victoria B.C., Los Angeles, CA and San Diego, CA (Fig 1A, B, Table S1). For site selection in California regions, we used data from the US National Oceanic and Atmospheric Administration (NOAA) Mussel Watch program (Dodder et al., 2014; Maruya, Dodder, Schaffner, et al., 2014; Maruya, Dodder, Weisberg, et al., 2014). Mussel Watch collects bivalve tissue and sediment samples across the US to identify areas with contaminants of emerging concern, trace metals and other contaminants. *Mytilus* mussels are used as the indicator species across the western US coastline because they reside in the intertidal environment, filter water for feeding and are widespread across this range. Mussel Watch has classified 87 intertidal sites across California based on surrounding land use (urban, mixed development, low development or agricultural), treatment plant proximity, and pollutant concentrations in *Mytilus* tissue and sediment (Dodder et al., 2014; Maruya, Dodder, Schaffner, et al., 2014, 2014). They found significant differences in pollutant concentrations between the different development categories, suggesting urban sites may have higher pollution exposure compared to the other development categories (Dodder et al., 2014; Maruya, Dodder, Schaffner, et al., 2014, 2014). For our California sampling, we designated sample sites as “nonurban” if they were closest to Mussel Watch “mixed development” and “urban” if they were closest to sites designated by Mussel Watch as “urban” (Fig S1). Our nonurban sites were on average 29km (range: 17.48km to 78.77km) away from an outflow site and our urban sites were on average 4.3km (range: 1.04km to 8.48km) from a wastewater outflow site. Within California, we sampled both intertidal and subtidal sites. The four sites in Victoria were chosen using similar classifications to Mussel Watch, considering surrounding land use and proximity to wastewater treatment plants for sampling sites. Victoria, B.C. provides a geographically distinct city in order to disentangle other environmental variables such as temperature, salinity and pH (Johnstone & Mantua, 2014) and all sites were intertidal. Urban sites were selected in close proximity to the McLoughlin Point Wastewater Treatment Plant, while nonurban sites were selected further west along the Salish Sea by the Strait of Juan de Fuca. One urban site was even colloquially named “garbage point” by locals due to the historical buildup of trash in the region (Clover Point in Victoria B.C., personal comm.).

**Figure 1:**
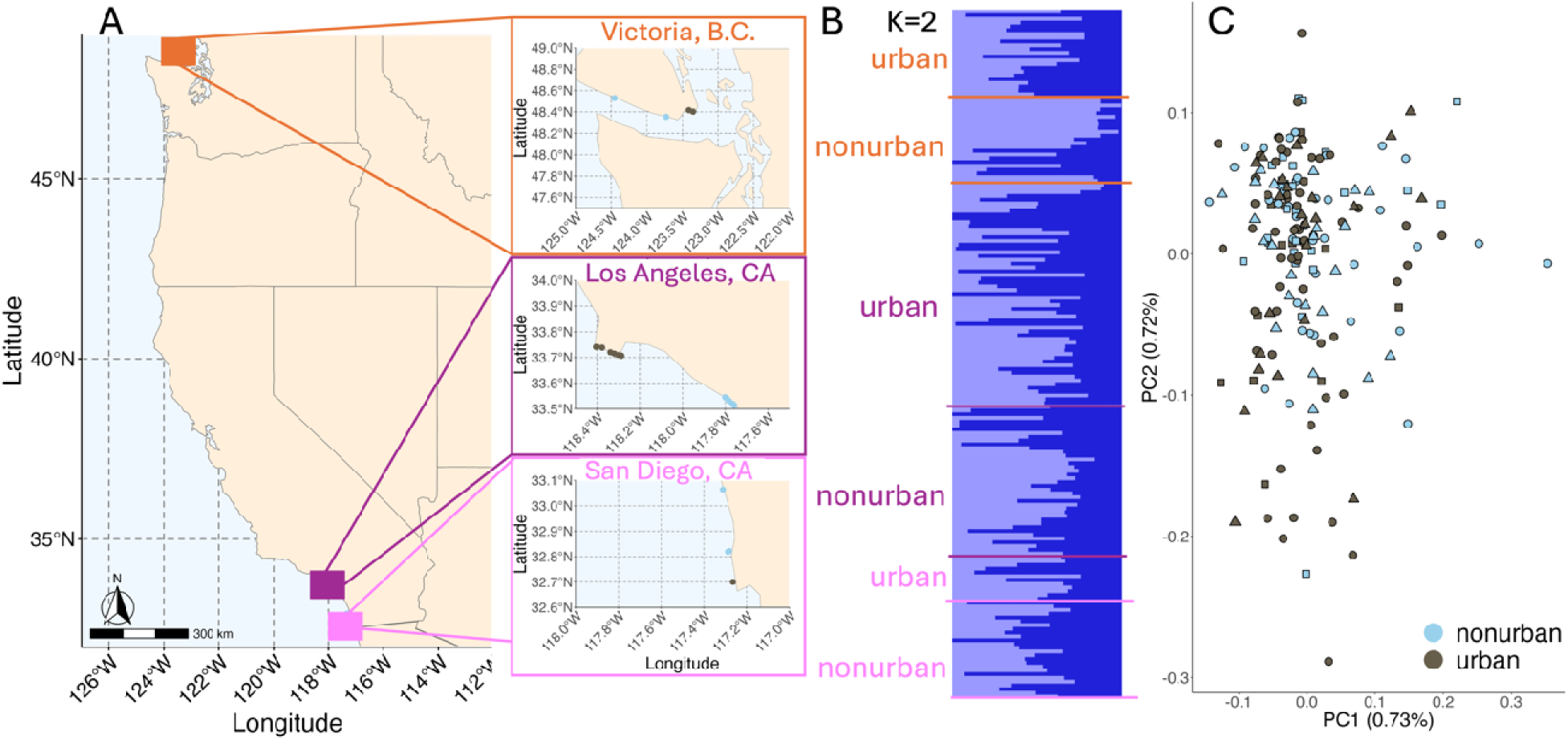
High gene flow in S. purpuratus sampled across three distinct coastal regions: Victoria, B.C., Los Angeles, CA and San Diego, CA. Both the population structure plot and the inset PCA were created using a thinned dataset of 19,299 SNPs to reduce linkage bias. (A) Overall map of sampling sites. Scale and orientation noted in the bottom left corner. (B) Each site zoomed in to show detailed sampling locations for the different regions: Victoria, Los Angeles and San Diego. Colors indicate urban (brown) or nonurban (light blue) sites. (C) The structure plot colors refer to the groups observed across the samples, with K=1 being the best fit, but we plotted K=2 for visualization purposes. (D) In the PCA, shapes correspond to the three regions, and color again corresponds to urban (brown) and nonurban (light blue) samples.

### Collections & Sample Processing

We collected urchin spine tissue from 209 Pacific purple sea urchins sampled across 19 sites. All sampled individuals in this study were adult *S. purpuratus* that were on average 7 cm in diameter, well above the size of juvenile *S. purpuratus* (Sonnenholzner et al., 2011), and given typical growth rate and low adult dispersal have likely been in the sampled region for 4-5 years. We collected at 11 sites across the LA region, with 6 urban (4 intertidal, 2 subtidal) and 5 nonurban (3 intertidal, 2 subtidal) sites. For San Diego, 4 sites were sampled, with 1 urban site (subtidal) and 3 nonurban sites (1 intertidal, 3 subtidal). Subtidal sites were also collected for Los Angeles and San Diego by California Conservation Genomics Project collaborators (Rumberger et al., 2025). For each site, we collected several spine tissue samples from 10-15 urchins (exact numbers reported in Table S1) and stored them in 95% ethanol for DNA extractions using a Qiagen blood and tissue DNA extraction kit. Permits were obtained through the California Department of Fish and Wildlife (SCP S-201570005-20183). Permits for Victoria field sites were obtained through the Fisheries and Oceans Canada Department (XR 19 2022).

### Sequencing & Bioinformatics Methods

Whole Genome Sequencing libraries were made using the protocol in (Nielsen et al., 2024), following the Nextera Lite protocol with similar modifications (Rowan et al., 2019). Whole-genome sequencing was done on a NovaSeq lane with 150bp paired end reads. Samples were sequenced at an average of 7.5x coverage. SnpArcher (v. 0.1), an automated snakemake (v. 8) pipeline using python (v.3.11.4) (Mölder et al., 2021), was used to process the data from fastq format to a single vcf (Mirchandani et al., 2024). Briefly, fastp (v.0.20.1) was used to quality check samples, bwa was used to align samples to the reference genome, samtools (v.1.11) was used to convert sam to bam files, GATK4 (v.4.1.8.0) was used to call haplotypes and estimate genotypes for samples in this pipeline and vcftools (v.0.1.16) was used to apply depth filters to exclude genotypes with too low or too high coverage (min-meanDP > 8, max-meanDP < 20). We conducted additional downstream filtering using vcftools to discard single nucleotide polymorphisms (SNPs) with minor allele frequency less than 0.05, SNPs with <70% of individuals genotyped, and genotypes with genotype quality <10.

### Population Structure

To investigate population structure, we used a subset of SNPs thinned by keeping 1 SNP per 25kb window to reduce linkage disequilibrium. *S. purpuratus* has been shown to have low linkage disequilibrium (Petak et al., 2023). We then used hierarchical clustering implemented in Tess3r (v.1.1.0) to identify population structure in our dataset. We estimated ancestry using K=1 through K=8 using 10 replicate runs for each K. Pophelper (v.2.3.1) was then used for visualization of Tess3r results. We also conducted a principal components analysis (PCA) of the thinned SNPs using SNPRelate (v. 1.38.1) (Zheng et al., 2012).

### Outlier Analyses

Genome scans were used to identify genomic signals of adaptation with the full SNP dataset. We implemented these methods to compare urban and nonurban sites within regions and also to compare across the three regions, investigating broad-scale signals of adaptation likely associated with latitude. Outlier SNPs were identified using OutFLANK (v.0.2) (Whitlock & Lotterhos, 2015). OutFLANK is an F_ST_ based analysis that identifies loci that are highly differentiated between groups compared to the null distribution defined by the genome-wide distribution and groups are specified *a priori*. Because we are particularly interested in directional selection, not balancing selection, we identified candidates for selection in the right-tail outliers. SNPs were considered significant if the false discovery rate corrected p-value (i.e. q-value) was less than 0.01. This was done for region, urban status, and tidal zone. We also used OutFLANK to contrast urban and nonurban sites in each region separately. To test whether signals of selection are shared across the three regions, we used a slightly more permissive q-threshold of 0.1 and tested for overlap across the region-specific outliers.

Genome scans, especially at the moderate sample sizes achievable in natural systems, are likely to miss loci of small to medium effect size (Regoli et al., 2006). One approach that can increase power is to integrate effects across multiple neighboring SNPs using a sliding window approach (Booker et al., 2023; Manel et al., 2016). We therefore combined local PCA (Li & Ralph, 2019) and linear models to create a novel framework to identify windows of SNPs differentiating urban and nonurban regions. First, to find an appropriate window size, we ran 8 trials using window sizes ranging from 30 to 100 SNPs and ran a local PCA for each window. Linkage disequilibrium has been shown to drop drastically at about 1,500 bp of genomic distance in *S. purpuratus* (Petak et al., 2023). We found that our SNP density was about 1 SNP per 3,700 bp, thus we were unconcerned about small window sizes capturing only linked SNPs. We found that smaller windows were prone to producing NAs rather than PC coordinates due to missing data, so we moved forward with a window size of 100 SNPs which had a low fraction of windows where PCs could not be calculated (0.048%). We then ran ANOVA tests to find associations between the first two PC coordinates calculated for each SNP window and the explanatory variables: city region (i.e. Los Angeles, San Diego, or Victoria), urban vs nonurban, and the interaction between the two. To determine a p-value that accounts for multiple testing, we ran 100 permutations with randomized city region and urban designations. In each randomized dataset, we ran an ANOVA to test for associations between each PC in each window and the randomized metadata. For each explanatory variable in our models, we found the 1st percentile p-value for each randomization and set the p-value threshold for each term to be the minimum of those values. The F-statistic threshold was obtained in a similar manner, by calculating the 0.99 quantile per randomization and then taking the maximum of those values. The empirical/observed p-value was considered significant if it was as extreme or more extreme than these thresholds.

### Detection of Polygenic Signals

We used *pcadapt* (v.4.4.0) (Luu et al., 2017) to identify polygenic signals of adaptation in our dataset. *Pcadapt* uses principle component (PC) loadings that don’t assume demography (e.g. population structure; (Zheng et al., 2012)) and performs well with populations that have high admixture (Silliman, 2019). However, pcadapt does not explicitly test for selection among *a priori* groups, it finds the largest sources of variation in the population without regard to experimental design. Note also that this PCA analysis is different from the PCA generated to assess population structure because we include all SNPs rather than just a thinned subset. To assess which of the multiple environmental variables present in our dataset were driving the variation between outlier SNPs, we conducted a redundancy analysis (RDA) (Capblancq & Forester, 2021; Lotterhos, 2023). We first trimmed the data by minor allele frequency (0.05) and then read in all environmental and sample predictors (sample ID, site ID, latitude, longitude, region, urbanization and tidal zone). We removed variables that were highly correlated with others and ran the RDA model with only region, urbanization and tidal zone. We then conducted backwards model selection with ordistep through the R package vegan (v. 2.7.1) similar to analysis done in Rumberger and Armstrong et al. (2025).

Complementary to the polygenic approach using RDA, we created a polygenic score by summing weighted effects across all SNPs (Simon et al., 2020). This approach has previously been used to predict polygenic phenotypes in medical and agricultural applications (Jayasinghe et al., 2024; Kullo et al., 2022; Ma & Zhou, 2021) and is gaining traction in ecological settings (Babin et al., 2017; Fuller et al., 2020). These methods sum effect sizes across all SNPs to create a single estimate of the response variable and can perform better than genome scans in modeling the genetic basis of phenotypes in natural systems (Fuller et al., 2020; Simon et al., 2020). We randomly selected training sets consisting of 60% of individuals and validated the model on the remaining 40%, and for each SNP we assessed the relationship between environment (urban or nonurban) and genotype. Beta coefficients were calculated using a latent factor mixed model (LFMM). We then used these beta coefficients to predict a polygenic score for each sample in the validation set, and repeated this test 100 times with different splits of the data, ensuring training and validation sets always had samples from each region and urban/nonurban sites. From each run, we used a t-test between predicted values (polygenic scores) for each sample in the validation set to determine whether urban and nonurban validation samples were statistically different. For visualization, we calculated the mean for each region/urban status group and plotted the distributions of those means across 100 runs.

To assess if our polygenic score test was actually predicting differences between urban and nonurban individuals, and not just separating individuals into two random groups by chance, we also ran a null model where the urban/nonurban labels were randomized. We followed the same methods above, running the test 100 times with different training (60% of the data) and validation (40% of the data) splits. We expect no difference in polygenic scores between randomly assigned groups, which provides confidence that real differences between urban and nonurban polygenic scores represent a true biological signal. We conducted two additional tests to validate our results. First, we conducted a separate analysis using a single coastal region as the training set and the remaining two as the validation sets for three total comparisons. Second, to ensure that results were not due to uneven sampling across regions, we ran a round of polygenic score tests using down-sampling to account for sample size variation between the three regions. We randomly sampled each population to the lowest sample size (n=38) 100 times. Although the reduction in sample size might result in reduced power, we expect that patterns should be qualitatively similar to those achieved in the full model if the signal is not due to differences in sample size across the three regions.

## Results

### Sequencing & Bioinformatics

For our 209 samples, we obtained an average of 14 Gb of raw sequence data per sample and identified 280,720,325 SNPs prior to filtering. Raw reads are available on NCBI (see data availability section). 26 samples were removed via downstream filtering due to low coverage (<3.2x). The removed samples included individuals from each region and both urban/nonurban individuals (Los Angeles: 8/7, San Diego: 3/4, Victoria: 2/2). This left 183 samples at an average of 8.6x coverage for the filtered dataset. After filtering, we identified 219,773 high quality SNPs across the *S. purpuratus* genome.

### Population Structure

We used a thinned set of 19,299 SNPs to analyze population structure, reducing effects of linkage disequilibrium. Clustering analysis in Tess3r (Caye et al., 2016) revealed the most likely scenario is a single ancestral lineage (K=1) and visual inspection of analyses run with K=1-6 clusters shows no geographical signal (Fig S2). Principal Components Analysis (PCA) on the thinned SNP dataset also shows no clustering by coastal region (Fig S3) or by urban status (Fig 1D).

### Outlier Analyses

Using the full set of 219,773 SNPs, we first conducted F_ST_ outlier scans across the entire genome and found no outliers significantly differing between city regions (Fig 2A). Somewhat surprisingly, F_ST_ among regions was lower than within region urban/nonurban comparisons (among region mean=0.0001, max=0.131). We did find loci that are highly differentiated between urban and nonurban environments within each coastal region and overall (Fig 2B-D, Fig S4). Genome-wide average F_ST_ between urban and nonurban sites were low in each of the three coastal regions, though higher than among regions (Victoria: 0.0012, Los Angeles: 0.0006, San Diego: 0.0008), and individual loci often had much higher differentiation (max F_ST_ for Victoria: 0.379, Los Angeles: 0.201, San Diego: 0.532). Analyzing all three regions jointly, we identified 165 SNP outliers associated with urban status (Fig S4). When we separated by region to assess outliers associated with urban status we found 17 outliers in Victoria, 741 outliers in San Diego and 2,271 outliers in Los Angeles (Fig 2B-D). There was little outlier overlap between regions; seven outliers were shared between Los Angeles and San Diego, and one was shared between Los Angeles and Victoria (Table 1).

**Figure 2:**
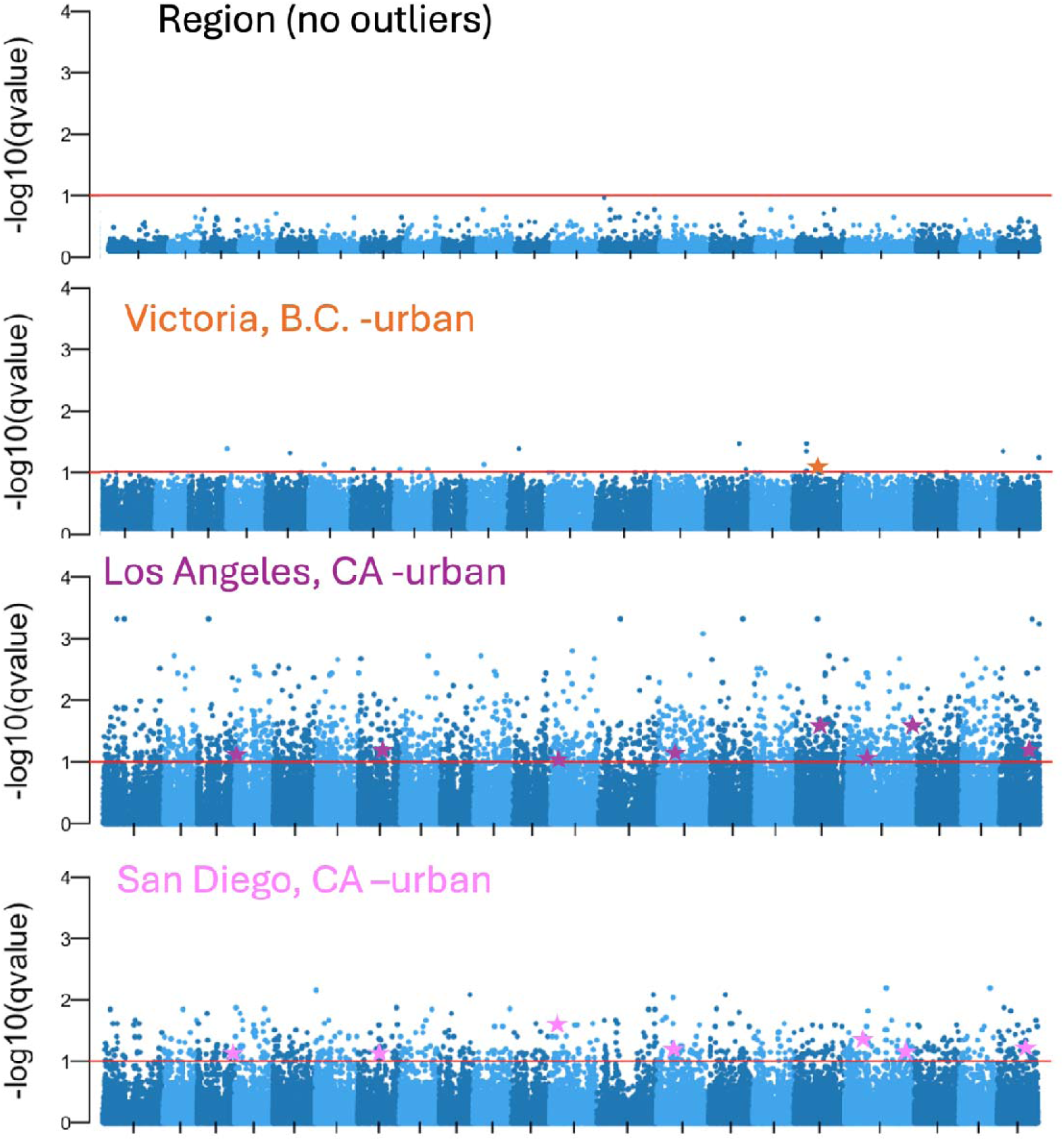
No genomic signals of selection associated with region, but signals of selection with urban status across three coastal regions. The x-axis of the Manhattan plots for OutFLANK (v.0.2) are the scaffolds in the *S. purpuratus* genome. The red line indicates that an outlier cutoff of p<0.1. A) There are no outliers that correspond with region for this dataset. B-D) Manhattan plots for urbanization across the three coastal regions. There are 17 outliers in Victoria (B), 2271 outliers in LA (C) and 670 outliers in San Diego (D) associated with urban status. The stars on each Manhattan plot indicate the shared outliers between the regions. Los Angeles and Victoria share 1 SNP outlier and Los Angeles and San Diego share 7 SNP outliers (Details on shared outliers in Table 1).

**Table 1:** Shared outlier genes identified between coastal regions. Outlier genes were identified via OutFLANK (v. 0.2) and shared genes between urban sites in the three coastal regions (Los Angeles, CA, San Diego, CA, and Victoria, B.C.) are reported below with function and gene type identified using gene ontology (GO). Columns include chromosome (scaffold), position, gene function from GO, gene type, transcript name and what regions this outlier was found in. Seven outliers were found to be shared between Los Angeles and San Diego, with one having no GO function identified, and one outlier was shared between Los Angeles and Victoria.

| Chromosome | Position | Function | Gene Type | Transcript Name | City Regions |
| --- | --- | --- | --- | --- | --- |
| NW_02214561<br>1 | 21363591 | coiled-coil domain-containing protein 34-like | protein-coding | XM_030998690.1 | Los Angeles & Victoria |
| NW_02214559<br>6 | 33274999 | caseinolytic peptidase B protein homolog | protein-coding | XM_030981235.1 | Los Angeles & San Diego |
| NW_02214560<br>0 | 19562203 | nucleolysin TIAR isoform X1 | protein-coding | XM_777843.5 | Los Angeles & San Diego |
| NW_02214560<br>5 | 5842857 | probable E3 ubiquitin-protein ligase HERC4 | protein-coding | XM_030992502.1 | Los Angeles & San Diego |
| NW_02214560<br>7 | 13648930 | NA | NA | NA | Los Angeles & San Diego |
| NW_02214561<br>2 | 16999057 | glycine receptor subunit alpha-3-like isoform X1 | protein-coding | XM_030972553.1 | Los Angeles & San Diego |
| NW_02214561<br>2 | 54981990 | protein DD3-3 | protein-coding | XM_796784.5 | Los Angeles & San Diego |
| NW_02214561<br>5 | 24968672 | ras-specific guanine nucleotide-releasing factor 2 isoform X1 | protein-coding | XM_011681667.2 | Los Angeles & San Diego |

From our sliding SNP window analysis, we identified 2,161 total windows. We tested for signals of selection associated with urban status or coastal region. We also tested for an interaction between urban status and coastal region, which would suggest that populations from different cities may be adapted to urban environments using different genetic mechanisms. Of the 2,090 usable windows (removing NA PCs), we found 53 outlier windows comparing urban vs. nonurban locations, 47 outlier windows associated with coastal region and 56 outlier windows with significant interaction between the urban and region variables (Fig 3A). Although the number of genomic windows with signals of selection associated with urbanization and coastal region were comparable, we did find that the effect size (F statistic) was larger for urban outliers (F-stat= 8.748) than for comparisons among coastal regions (F-stat=5.837) or the interaction term (F-stat=6.038) (Fig 3B-C), supporting our SNP-level findings differentiating urban and nonurban sites. However, the window-based analysis uncovered signals of selection absent from the SNP-based analysis; only five outlier SNPs from our combined analysis fell in significant windows (Fisher’s exact test p=0.603).

**Figure 3:**
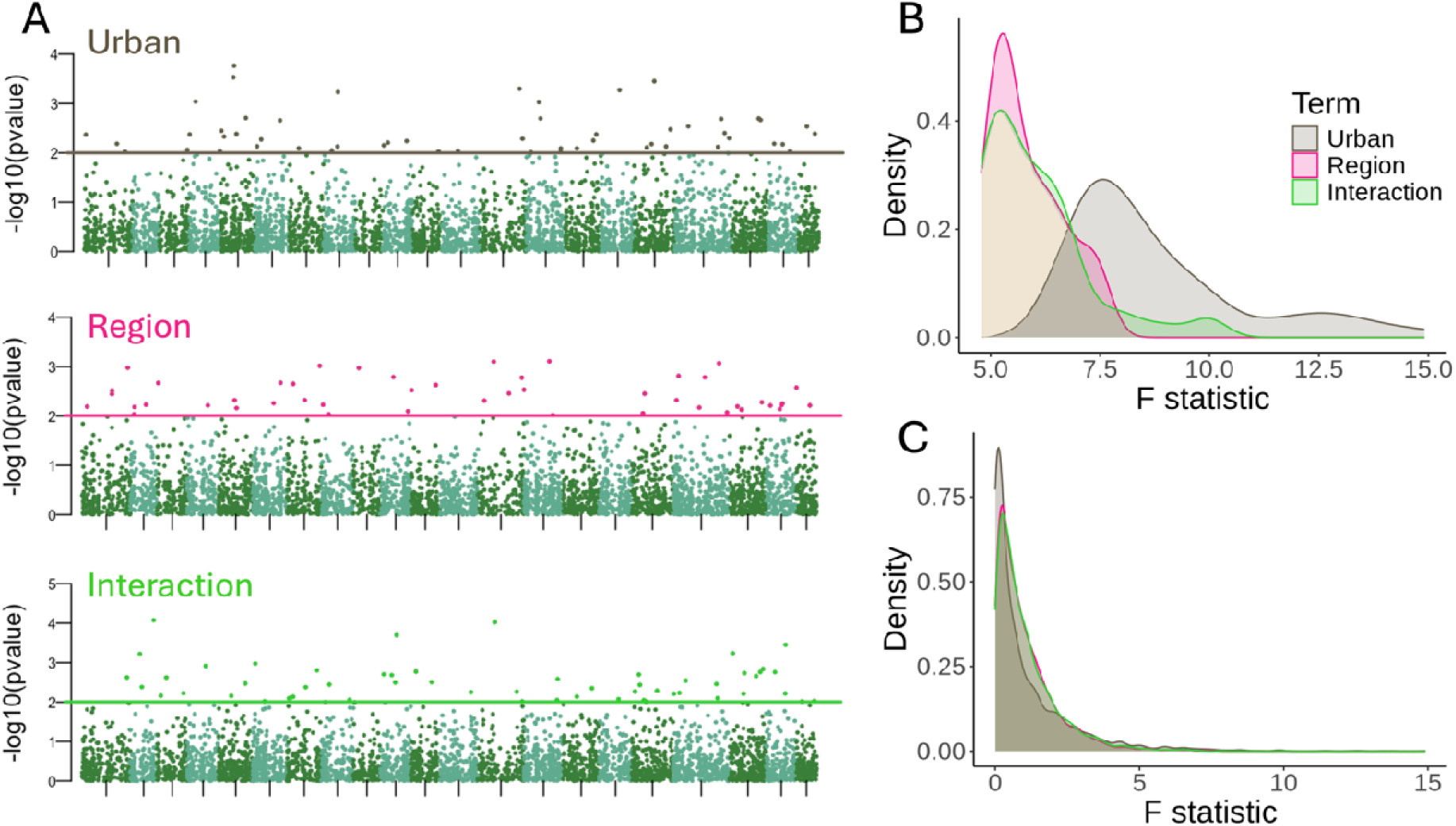
Sliding window analysis identifies signals of selection associated with urban status and coastal region. Colors represent different terms: urban (brown), coastal region (pink) and the interaction between the two (green). Windows of 100 SNPs were used to test significant associations with urban, coastal region or the interaction. (A) Manhattan plots of each term of interest and the SNP window outliers associated. The points below the threshold line on each plot are nonsignificant SNP windows while points above the line represent significant SNP windows. P-value thresholds are determined by randomization. For urban status (brown), the p-threshold was at 0.0098 and 53 outliers were identified. For the coastal region (pink), the p-threshold was at 0.0098 and 47 outliers were identified. Finally for the interaction (green), the p-threshold was at 0.0099 and 56 outliers were identified. (B-C) F-statistic distributions of P-value outliers for each group for significant (B) and nonsignificant (C) SNP windows. (B) Mean effect sizes were larger for urban significant SNP windows compared to coastal region or the interaction significant SNP windows. Note the x-axis begins at 5, rather than 0, for visualization purposes. (C) This effect is not seen in the null distribution. Mean effect sizes of non-significant SNP windows did not differ across region, urban or the interaction, thus this is unlikely to be an artifact of degrees of freedom differences between our terms (region=3 while urban=2).

### Detection of Polygenic Signals

Using multiple polygenic approaches, including *pcadapt*, a redundancy analysis (RDA) and polygenic scores, we find distinct differentiation between urban and nonurban samples across the three regions. For our RDA, we reduced our full predictor set to the uncorrelated predictors of latitude, urban status and tidal zone. Using backwards model selection we found that urban status (F_1,238_= 1.046 p=0.023) and tidal zone (F_1,241_= 1.062 p=0.005) were significant, but latitude was not (F_1,230_= 1.012 p=0.124) (Table S1; Fig 4). We then conducted several principal component analyses (PCA) using *pcadapt* on the full SNP set to identify differentiation by region, urban status and tidal zone. There were no significant PC axes for region (Fig 5A-C). Several PCs differentiated across tidal zone (Fig 5G-I), including PC2 (A, t-value= 2.293, p=0.023), PC3 (B, t-value=-2.908, p=0.0041) and PC4 (B, t-value=3.588, p<0.001), consistent with previous findings of selection between intertidal and subtidal sites (Rumberger et al., 2025). For urban status, we identified several PCs of interest, including PC3 (t-value=2.941, p=0.0037), PC5 (t-value= 4.997, p<0.001) and PC6 (t-value= -4.586, p<0.001). PC1-PC6 for urban status are shown together in the supplement (Fig S5). Focusing on our most significant axes of variation (PC5 and 6, Fig 6A), we also found that differentiation between urban and nonurban environments on *pcadapt* PCs were not driven by a single city region; analysis of PC loadings showed that on these PCs, urban and nonurban samples were differentiated across either two (Fig 6B) or all three city regions (Fig 6C).

**Figure 4.**
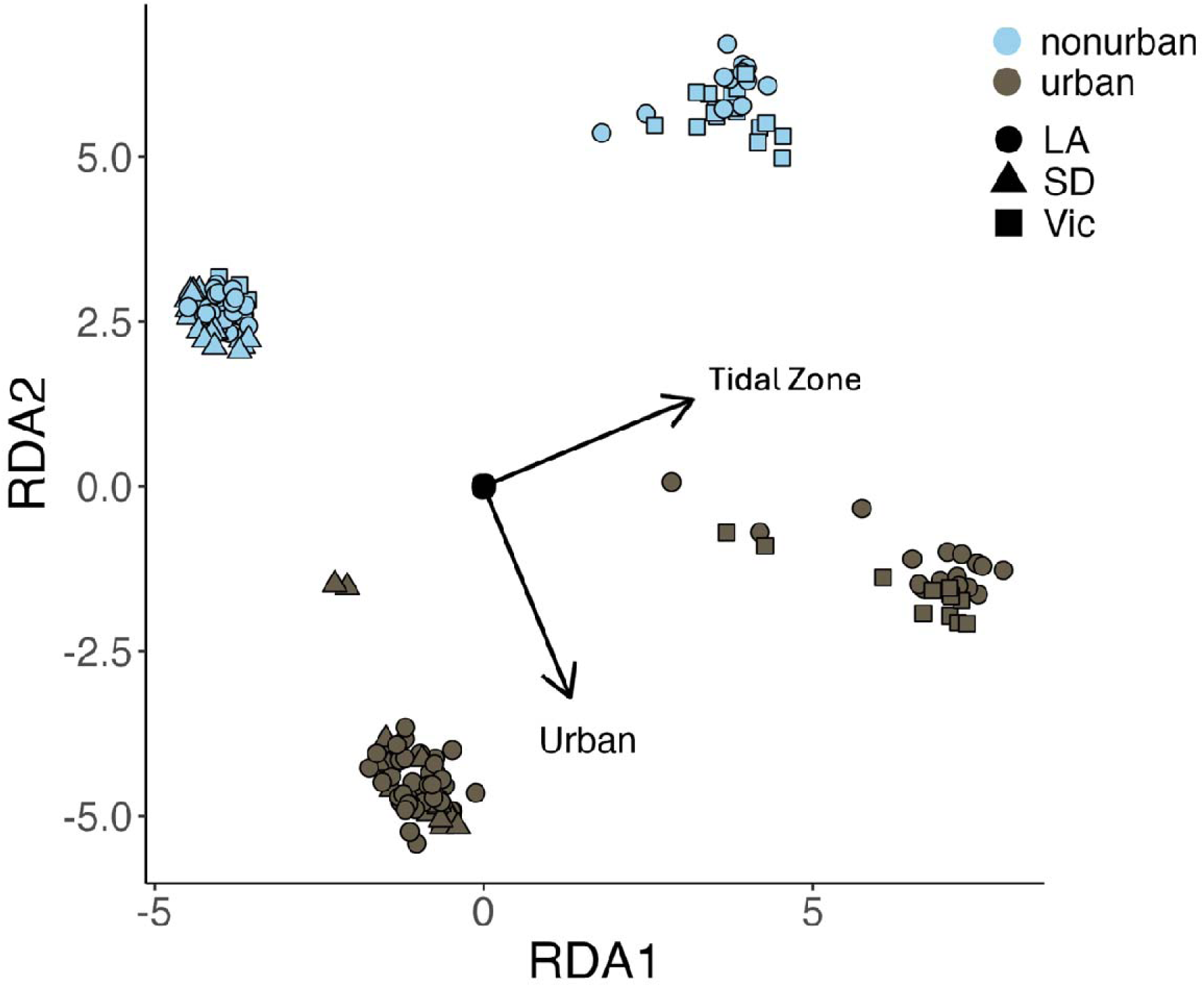
Redundancy Analysis shows signals for urban status and tidal zone but not latitude. Colors distinguish urban (brown) and nonurban (light blue) samples, while shapes denote coastal region (Los Angeles, San Diego and Victoria). Only urban status and tidal zone were significant for model prediction, while latitude was not. RDA1 showed the spread of data across the intertidal to subtidal gradient, while RDA2 showed the separation of urban and nonurban samples.

**Figure 5:**
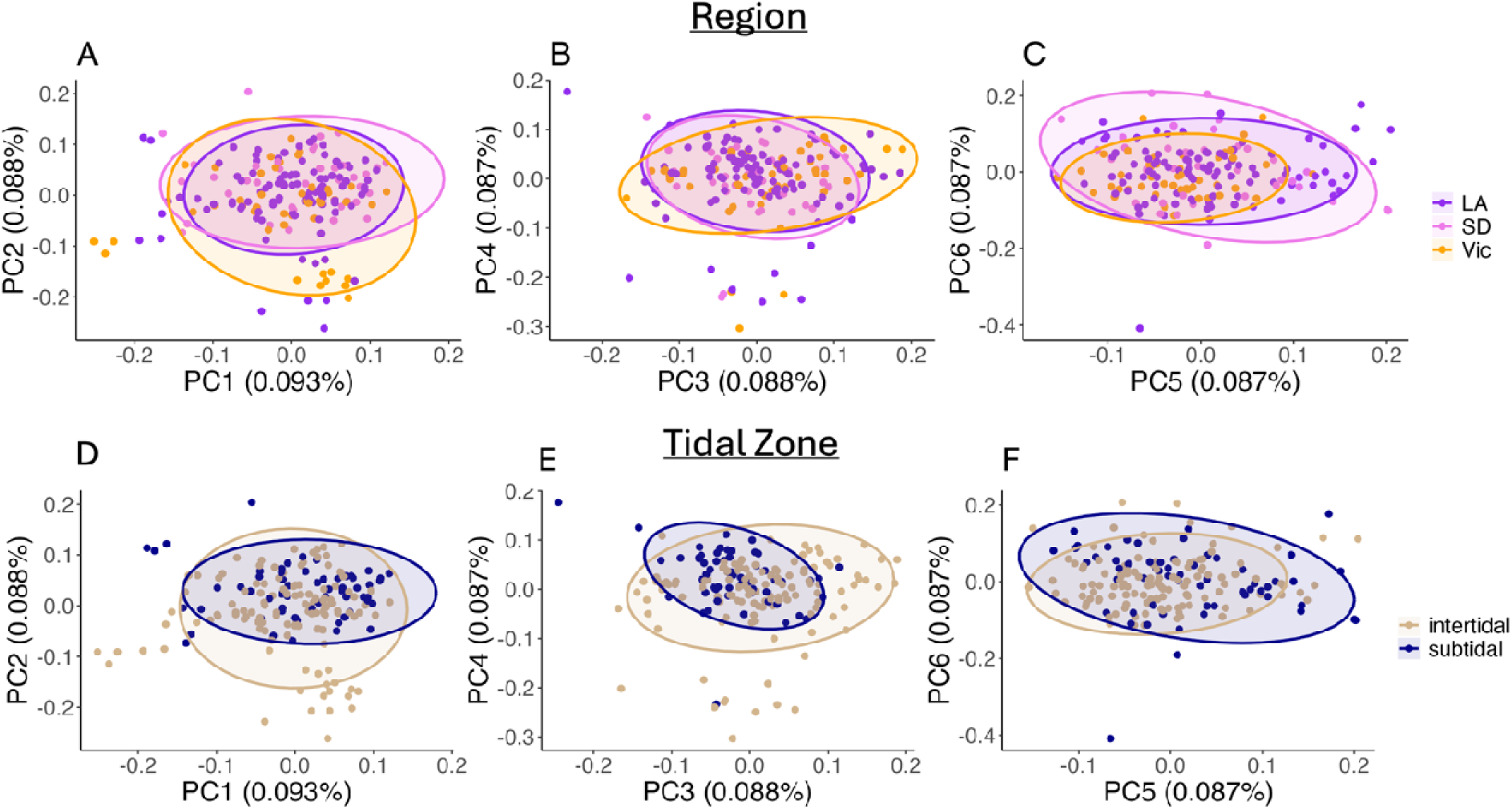
PC axes show no differentiation, between region, but evidence of variation between tidal zones. Principle components (PCs) were identified using *pcadapt* (v. 4.4.0). PC1-PC6 shown here for regio (A-C) and tidal zone (D-F). (A-C) No PC axes show significant variation between our three coastal regions. Each region is a distinct color: Los Angeles, CA (LA) in purple, San Diego, CA (SD) in pink and Victoria, B.C. (Vic) in orange. (D-F) Variation at several PC axes observed between samples across different tidal zones. Colors distinguish intertidal (tan) and subtidal (navy blue) samples. We found that tidal zone drove variation along several PC axes, included PC2 (E, t-value= 2.293, p=0.023) and PC3 (F, t-value=-2.908, p=0.0041) and PC4 (F, t-value=3.588, p<0.001). This was not unexpected given previous literature highlighting SNP outliers between subtidal and intertidal *S. purpuratus* populations across California (Rumberger et al., 2025).

**Figure 6:**
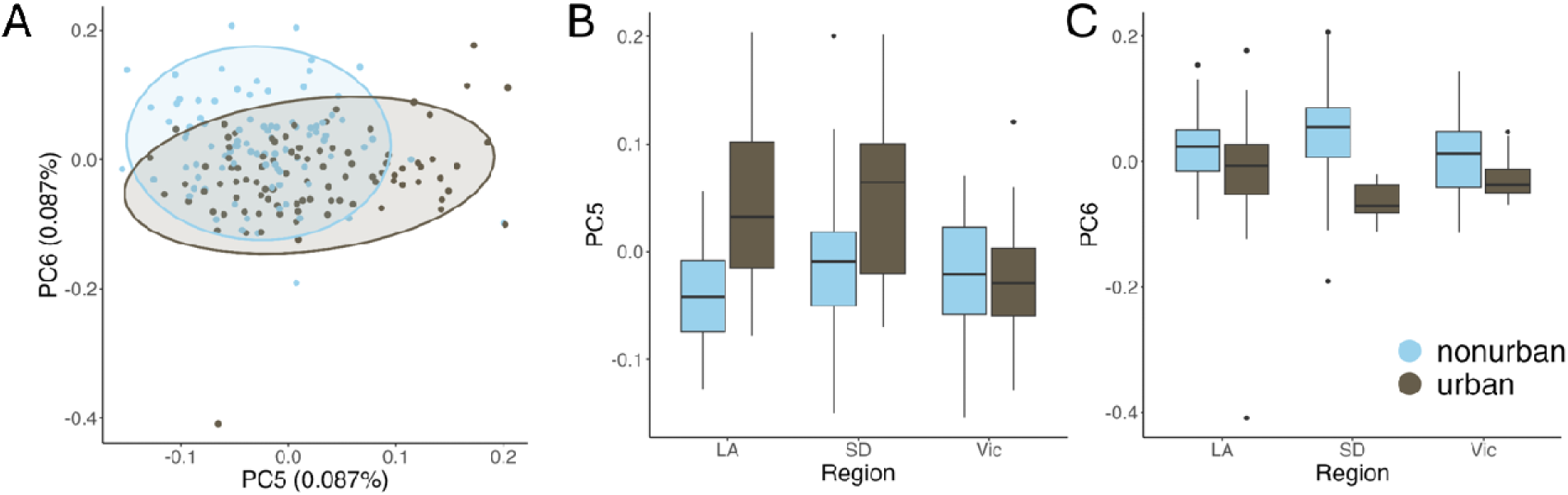
Differentiation between urban and nonurban samples not driven by one coastal region. Principle components (PCs) were identified using *pcadapt* (v. 4.4.0). Urban sites are shown in brown, while nonurban sites are shown in light blue. (A) PC5 and PC6 showed significant variation between urban and nonurban samples. (B-C) This differentiation was not driven by one coastal region, with urban and nonurban samples separating in Los Angeles (LA), San Diego (SD) and Victoria (Vic) as well for both PC5 (B) and PC6 (C) respectively.

For our polygenic score analysis, 90/100 runs yielded significantly different predicted scores between urban and nonurban samples in the validation groups (Fig 7A). When urban status was randomly assigned, we saw no statistical difference between groups (Fig 7B). To further interrogate parallel signatures using this approach, we conducted a separate analysis using a single city region as the training set and the remaining two as the validation sets for three total comparisons (Fig S6). Here we found differences in the predictive ability depending on the population used for training. A model trained on San Diego populations differentiated between urban and nonurban sites (two-way ANOVA, F_1,1_= 6.845 p=0.010) but more strongly in Victoria (two-way ANOVA interaction, F_1,1_= 5.493, p=0.020) (Fig S6B). Reciprocally, the model trained on Victoria populations accurately predicted urban status in San Diego, but not in Los Angeles (Fig S6C, two-way ANOVA interaction, F_1,1_= 9.699, p=0.002). The model trained on Los Angeles did not perform well, with no differences between predicted scores for urban and nonurban samples in either Victoria or San Diego (two-way ANOVA, F_1,1_= 0.341, p=0.561). Lastly our down-sampled polygenic score test still showed consistent patterns differentiating urban and nonurban samples across each city region, with 71/100 runs yielding significantly different predicted scores between urban and nonurban samples (Fig S7).

**Figure 7:**
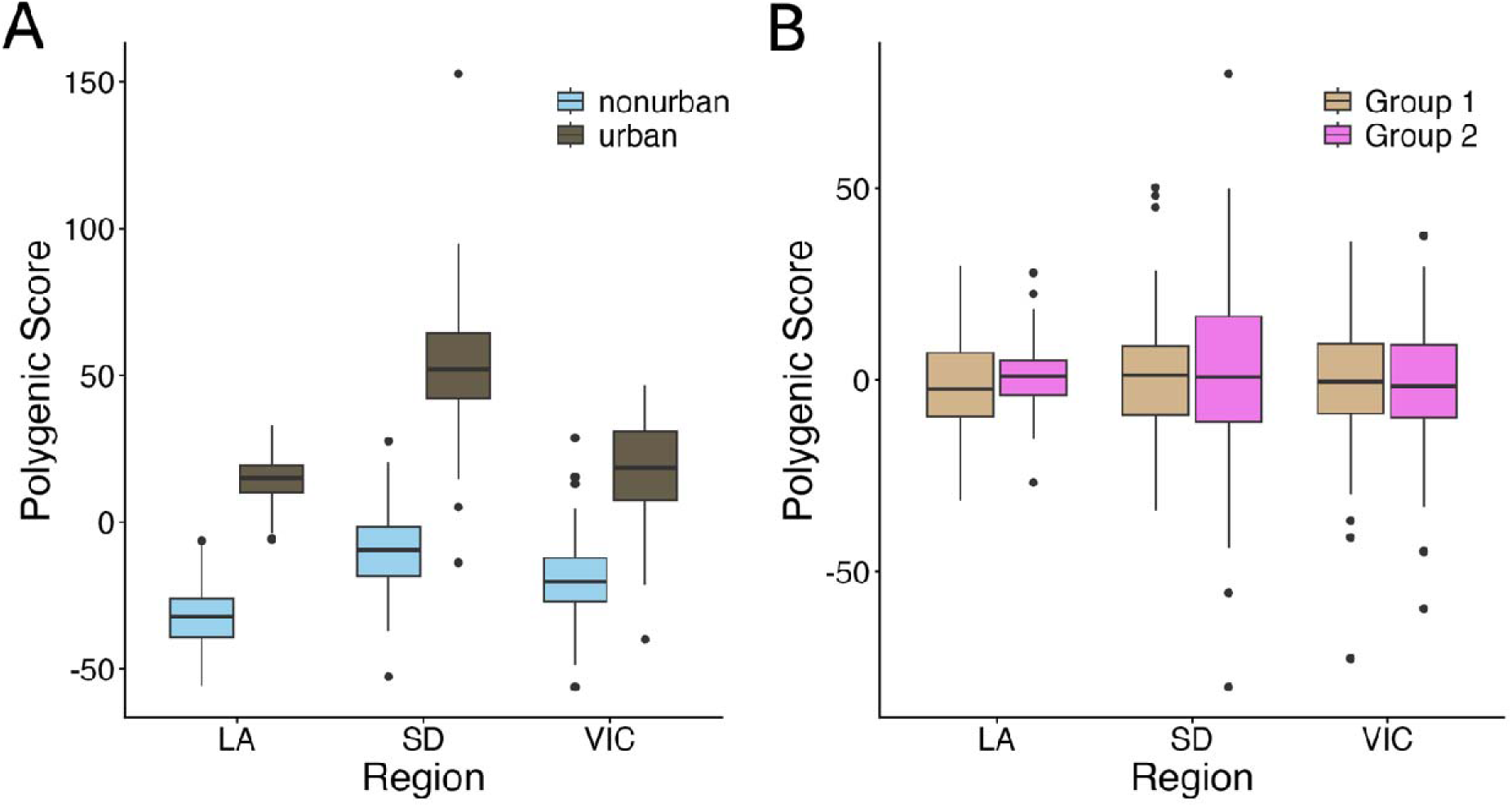
Polygenic score test highlights significant differences between urban and nonurban samples was not random. To validate that the separation of urban and nonurban samples was nonrandom, we implemented a polygenic score test (PGS). The x-axis separates by our three regions (Los Angeles (LA), San Diego (SD) and Victoria, B.C. (VIC)), while the y-axis highlights the polygenic score. A) PGS for urban compared to nonurban samples. Urban sites are shown in brown, while nonurban sites are shown in light blue. After 100 runs, we found that 90/100 runs identified significant differences between urban and nonurban samples (p<0.05). We also assessed if each coastal region could predict urban/nonurban variation in the other two coastal regions (Fig S6) and the impact of sample size on our polygenic score test (Fig S7). B) In our null model PGS, we randomized the labels for urban and nonurban sites into either group 1 (tan) or group 2 (pink) to assess if the groups were separating by random chance. This null model showed no ability for the polygenic score to differentiate samples. After 100 runs, there is no significant differentiation from either group, supporting that the separation of our data into urban and nonurban groups in 7A is unlikely to be random chance.

## Discussion

This study not only adds to the growing literature on urban evolution in marine systems (Fusco et al., 2021; Park et al., 2025; Todd et al., 2019; Whitehead et al., 2017), but leverages genomic techniques to disentangle how urbanization shapes contemporary evolutionary processes in real time (Carlen et al., 2026). Our study uses whole genome sequencing to demonstrate evidence supporting urban adaptation in a highly dispersing marine invertebrate, the Pacific purple sea urchin (*Strongylocentrotus purpuratus).* While there was no evidence of population structure across the >2000km range, we identified genetic variants that differentiated between urban and nonurban sites within each city region. Additionally, despite little overlap in SNP outliers, we did find strong evidence of parallel signals of adaptation through polygenic approaches. Overall, our results show that even with high gene flow, there can still be polygenic and parallel signals of adaptation in marine invertebrate species in response to novel microgeographic variation.

### Lack of population structure across large geographic range

Many marine species, especially those with a planktonic larval phases, have large ranges and experience high gene flow across their species range (Hedgecock, 1986; Hellberg, 2009; Marko & Hart, 2017). For example, a recent study on red abalone (*Haliotis rufescens*) found no evidence of population structure and high shared genetic diversity between populations spanning from Oregon (USA) to Baja California, Mexico (Griffiths et al., 2025). Even with sedentary adult phases, the movement of larvae with ocean currents can result in little to no population structure across populations. *S. purpuratus* has a planktonic larval phase that lasts from 30–86 days (Strathmann, 1978), which allows some offspring to settle extremely far from their parents. Although our study extends much farther north than prior sampling efforts for *S. purpuratus*, we still find no evidence of barriers to gene flow or isolation by distance across >2000km (Fig 1C, D). Together with other studies, these findings support one large well-mixed population of *S. purpuratus*.

### Potential drivers of urban adaptation

Despite extremely high gene flow across the range, we find genomic signals of selection associated with the urban environment in each of the three coastal regions. Previous studies in marine invertebrate species have suggested that balanced polymorphisms in marine systems can lead to adaptation over small spatial scales even with high gene flow (Rumberger et al., 2025; Sanford & Kelly, 2011; Tepolt et al., 2022). With high gene flow resulting in homogenization between populations, we might also expect populations to share a substantial amount of standing genetic variation, potentially leading to the reuse of shared alleles in small-scale adaptation. Despite this, we find very little overlap in SNPs associated with urban status between our three sampled regions. We find only eight overlapping between the three regions (Table 1). Of these SNPs, two were found to be associated with oxidative stress, which is a generalized stress response with known ties to temperature and other stressors (Granados-Cifuentes et al., 2013; Selmoni et al., 2021). Several studies have found oxidative stress responses associated with urbanization (Herrera-Dueñas et al., 2014, 2017; Isaksson, 2015, 2020; Regoli et al., 2006), suggesting a potentially conserved mechanism that merits deeper investigation.

Our sites were selected using a combination of land use data, locations of wastewater outflows, and pollutant data from Mussel Watch, yielding a broad urban versus nonurban comparison. However, our findings alone cannot disentangle the many potential mechanisms of selection to marine urban environments. Urban impacts on marine environments are varied, including stormwater and sewage discharge, hardened shorelines, and increased human use (Alter et al., 2021). Wastewater outflow has also been shown to result in decreased pH, increased water temperature and decreased salinity (Gierach et al., 2017; Lin et al., 2025; Washburn et al., 1992), which on their own have been shown to affect physiology and evolution in marine species (Gleason & Burton, 2015; González Durán et al., 2018; Griffiths et al., 2021; Hoegh-Guldberg et al., 2007; Hughes et al., 2017; Kültz, 2015; Przeslawski et al., 2015; Wong & Hofmann, 2020). While our study doesn’t focus on a specific urban stressor, some chemicals have been shown to be in much higher concentrations in urban wastewater, including alkylphenols which were identified by the Mussel Watch program to be in higher concentrations in urban areas along the California coastline (Dodder et al., 2014, 2014; Swam et al., 2024). Adaptation to these novel urban environments can be rapid. For example, in response to polychlorinated biphenyl (PCB) exposure, which is highly correlated with Superfund sites, Atlantic killifish have been shown to rapidly adapt with little phenotypic abnormalities but large adaptive shifts in their AHR pathways to mitigate stress responses (Whitehead et al., 2017). Experiments exposing urban and nonurban populations to pollutants associated with urban environments will be required to begin to disentangle the mechanisms of adaptation. Another potential stressor associated within many of our urban sites is harvesting pressure. Most of our nonurban sites were in marine protected areas (MPAs) where collection is prohibited, while urchins could be harvested with a fishing permit at many of the urban sites (sites labeled in Table S1). Fishing pressure has been shown to result in decreased genomic variation and result in selection in a variety of species (Helmerson et al., 2025; Mendoza Portillo et al., 2023; Nielsen et al., 2024; Reid et al., 2023; Sadler et al., 2024). This selection can occur at a rapid timescale resulting in population-level differences. While our results show signals of selection in response to urbanization, further studies are needed to determine which of the many urban stressors contribute most to selection.

### Genomic signals of adaptation over small, but not large, spatial scales

Intuitively, we would have expected Los Angeles and San Diego to have stronger parallel signatures as they are more likely to share standing genetic variation due to geographic proximity (Rennison et al., 2020). One possible explanation for a lack of a large number of shared urban outliers between our three coastal regions could be that the urban environments themselves represent very different selective environments. Los Angeles is a much bigger city, with a population of 3.8 million while San Diego and Victoria are at 1.4 million and 398,000 respectively. The cities differ in their development history, wastewater treatment methods, shipping port activity, etc, which could be driving genomic differences in nonurban and urban samples. This type of heterogeneity across cities has been highlighted in several studies (Dant et al., 2025; Santangelo, Roux, et al., 2022). Urban-associated F_ST_ outliers were smaller in magnitude in Los Angeles than the other two cities (max F_ST_ for Victoria: 0.379, Los Angeles: 0.201, San Diego: 0.532), perhaps reflecting a more complex landscape of selection than our binary urban/nonurban classification captures. Despite this, our analysis of the full dataset did find a polygenic model that captured genetic differentiation in response to urban environments across all three coastal regions.

We also find little overlap between our window and SNP-based outlier analyses. Only five outlier SNPs from our combined analysis fell in significant windows (Fisher’s exact test p=0.603), suggesting that our significant windows are likely made up of linked, small effect SNPs associated with urban status. The genomic windows significantly associated with urban status may reflect parallel signals that were not apparent in our individual SNP-level analysis. Because window-based analyses integrate across multiple SNPs across a genomic region, they can have increased power to pick up smaller effect size signals (Booker et al., 2023; Hoban et al., 2016). For example, a study in cattle found unique genes of interest using window-based methods that were undetected with standard SNP-based outlier methods (Braz et al., 2019). Additionally, window-based approaches can reduce the noise associated with linked SNPs (Moreira & Smith, 2023; Shi et al., 2023). Alternatively, the individual SNP-level outlier analysis may pick up SNPs that are overshadowed in the window-based analysis. SNP windows may include neutral and adaptive SNPs, which can result in a weaker signal from that window (Chaturvedi et al., 2025). While the sliding window approach is still novel in the field (Otte, 2024), both methods highlight important signals of adaptation and pick up on different metrics of diversification.

In contrast to signals of selection across urbanization gradients, which span small spatial scales, we see little evidence of selection across latitude. Our samples span 2000km, 16 degrees of latitude, 9 degrees Celsius of mean temperature, and a plethora of other environmental differences. Despite this, latitude was not significant in the RDA model, suggesting no strong signatures of selection across this broad environment gradient. PC axes from *pcadapt* were not explained by differences among regions and we found no outliers in SNP-level genome scans comparing regions. In our sliding window analysis, some windows were significantly different among coastal regions, but the effect size was lower than that comparing urban to nonurban sites. Previous studies have found evidence of selection associated with pH in populations spanning Oregon to San Diego (Pespeni et al., 2012, 2012, 2013; Pespeni & Palumbi, 2013) and weak but significant evidence of selection across the range of sea surface temperatures in California (Rumberger et al., 2025). Still, our results suggest that long-term selection to broad-scale climate gradients is perhaps not the predominant driver of genetic variation in this system.

This counterintuitive result of signals for selection at small, but not large spatial scales was also found in our previous work showing stronger support for selection across tidal zone than across climate gradients (Rumberger et al., 2025). Here, our RDA on independent samples also support this finding, with tidal zone (F_1,241_= 1.062 p=0.005), significantly contributing to the distribution of genetic variation and *pcadapt* PCs diverging across tidal zone. Selection over the small spatial scales are likely maintained through post-settlement mortality each generation (Levene, 1953; Pespeni & Palumbi, 2013; Thomas et al., 2018). This phenomenon has been observed in vertical zonation of barnacles along the intertidal zone (Schmidt et al., 2000) and in mussels across an estuarine gradient (Koehn et al., 1976). Because gene flow is so high in the *S. purpuratus* system, perhaps selection at smaller spatial scales is no less likely than at larger spatial scales, as both are derived from standing genetic variation in a single large population. In this case of microgeographic adaptation (Richardson et al. 2014), balanced polymorphisms are maintained across the species range which can then be reused in each city region. This can also increase the probability of polygenic parallelism (Rennison et al., 2020; Yeaman, 2015) if very high gene flow allows populations within each coastal region access the same putatively adaptive alleles.

### Parallel polygenic signatures of urban adaptation

Although there was little overlap in SNP-level F_ST_ outliers among regions, we find evidence for parallelism with our three polygenic methods. Because most complex traits are likely to be polygenic, approaches that more closely model this genetic architecture are increasing in use for understanding adaptation and trait evolution in natural populations (Gagnaire & Gaggiotti, 2016; Lotterhos, 2019). Genome scans including genome wide association studies (GWAS) and genotype environment associations (GEA) have, in many cases, uncovered large effect loci or even genomic structural variation underlying adaptation and the expression of ecologically important traits (Barrett et al., 2019; Johannesson et al., 2017; Vollmer et al., 2023). However, these methods are underpowered for quantitative traits, likely leading to a high rate of false negatives in natural systems (Hoban et al., 2016). In many cases, polygenic approaches can yield insight where genome scans fail. For example, Fuller et al. (2020) found no SNPs significantly associated with bleaching phenotype, but polygenic scores increased the power to predict bleaching. Similarly, Laporte et al. (2016) used polygenic approaches to model urban adaptation in American and European eels, uncovering signals of adaptation in sterol regulation pathways in response to pollution. Because polygenic approaches are explicitly predictive, they can easily be used to test for parallelism even when there is no overlap in SNP-level candidates for selection. In a study of eelgrass in the eastern Pacific, the authors found no shared SNPs responding to independent temperature gradients, but polygenic scores were able to predict temperature across both populations (Schiebelhut et al., 2024). With increasing studies of genomic data in natural systems, polygenic adaptation seems to be common across the tree of life and approaches that appropriately model polygenic selection are likely to yield novel insights (Csilléry et al., 2018; Forsberg et al., 2025; Gallardo-Hidalgo et al., 2025; Hämälä et al., 2020; Laporte et al., 2016; Moran et al., 2024; Pinseel et al., 2025; Rey et al., 2020; Rose et al., 2018; Yengo et al., 2022).

## Conclusion

Together, our results support parallel polygenic signals of selection across urbanization gradients in *S. purpuratus*, a broadly distributed coastal invertebrate. This is despite extremely high gene flow and a lack of latitudinal signatures of selection. We add to a growing literature suggesting that novel urban environments can result in selection on contemporary timescales (Santangelo, Ness, et al., 2022; Whitehead et al., 2017). Although there was no individual SNP that was “the urban SNP” for all three regions, we show that a lack of overlap at large-effect loci does not necessarily mean that genetic mechanisms of adaptation are independent; polygenic models find range-wide parallel microgeographic signals of urban adaptation. Increased use of polygenic approaches may uncover parallelism that is overlooked at the level of SNPs alone.

## Supporting information

Supplemental Tables 2-4 GO Terms

## General

Thank you to all of the fieldwork assistants who collected *S. purpuratus* spine tissue samples: Rob Dellinger, Mackenzie Kawahara, Kathryn Sutherland, Benjamin Lee, Camille Rumberger, Ed Parnell, Jason Toy, Zoe Scholtz and Adam and Jenesa Wall. Thank you to K. Lee for the urban color theme used throughout this paper. Thank you to Andrew Whitehead, Eric Sanford, Northeastern’s Popgen Fika group and all anonymous reviewers for providing feedback on this manuscript.

## Author contributions

Conceptualization: MLA, RAB. Methodology: MLA, RH, BC, KLE. Investigation: MLA, RAB, KLE. Visualization: MLA, RAB. Funding acquisition: MLA, RAB. Project administration: MLA. Supervision: RAB. Writing – original draft: MLA. Writing – review & editing: MLA, RH, BC, RAB, KLE

## Data and Materials Availability

Raw reads are available on NCBI accession: PRJNA1317549, vcf file on dryad: https://doi.org/10.5061/dryad.4xgxd25ph and all other scripts and data on github: https://github.com/mlarmstrong/urbanurchins

