## Supplemental Figures for "Parallel polygenic urban adaptation despite high gene flow in a coastal marine invertebrate"

**Supp Table 1:** Table of sample sites. This includes exact site latitude/longitude and what region they correspond to: Los Angeles (LA), San Diego (SD) or Victoria, B.C. (Vic). The table also includes site ID, tidal zone (intertidal or subtidal), whether the site was a marine protected area (MPA), the number of adult sea urchins sampled per site (n), and who collected the samples by last name.

| Site | Latitude | Longitude | region | Urban | Site_ID | Tidal Zone | California  MPAs | sample size (n) | Collector |
| --- | --- | --- | --- | --- | --- | --- | --- | --- | --- |
| Botanical Beach | 48.52918 | -124.4525 | Vic | nonurban | Bot | intertidal | N/A | 15 | Armstrong |
| Macaulay Point, Victoria BC | 48.41659 | -123.40953 | Vic | urban | Mac | intertidal | N/A | 10 | Armstrong |
| Clover Point, Victoria BC | 48.40281 | -123.35042 | Vic | urban | Clov | intertidal | N/A | 15 | Armstrong |
| Whiffin Spit, Sooke BC | 48.35431 | -123.72819 | Vic | nonurban | Whif | intertidal | N/A | 10 | Armstrong |
| Pelican Cove | 33.7409035 | -118.40399 | LA | urban | Pel | subtidal | Abalone Cove SMCA | 6 | Wall |
| Portuguese point | 33.73837 | -118.37577 | LA | urban | Port | intertidal | Abalone Cove SMCA | 10 | Armstrong/Lee |
| KOU Rock | 33.7203 | -118.33722 | LA | urban | KOU | subtidal | N/A | 12 | Scholtz |
| White Point, San Pedro | 33.71497 | -118.31871 | LA | urban | Wh | intertidal | N/A | 10 | Armstrong/Dellinger |
| Wilders Addition Park, San Pedro | 33.7096 | -118.29956 | LA | urban | WAP | intertidal | N/A | 10 | Armstrong/Kawahara/Dellinger |
| Cabrillo Beach, San Pedro | 33.70716 | -118.28562 | LA | urban | Cab | intertidal | N/A | 19 | Armstrong/Kawahara/Dellinger |
| Twin Points, Laguna Beach | 33.54529 | -117.80068 | LA | nonurban | Twp | intertidal | Laguna Beach SMR | 10 | Armstrong/Lee |
| Recreation Point | 33.54119 | -117.79276 | LA | nonurban | Rec | subtidal | Laguna Beach SMR | 5 | Toy |
| Laguna Beach | 33.52987 | -117.78093 | LA | nonurban | Lag | subtidal | Laguna Beach SMR | 12 | Williams |
| Sugarloaf Point, Orange County | 33.5197 | -117.76448 | LA | nonurban | Sug | intertidal | Laguna Beach SMR | 10 | Armstrong |
| Treasure Island Park, Orange County | 33.51372 | -117.75829 | LA | nonurban | Trp | intertidal | Laguna Beach SMR | 10 | Armstrong/Sutherland |
| Leucadia | 33.06373 | -117.31094 | SD | nonurban | Leu | subtidal | N/A | 5 | Scholtz |
| Big Rock Reef, San Diego | 32.8223 | -117.28095 | SD | nonurban | Brr | intertidal | South La Jolla SMR | 10 | Armstrong |
| La Jolla, San Diego | 32.8196 | -117.2871 | SD | nonurban | Bir | subtidal | South La Jolla SMR | 10 | Parnell |
| Point Loma, San Diego | 32.7007 | -117.2638 | SD | urban | Lom | subtidal | Cabrillo SMR | 15 | Parnell |

**Supp Table 2:** 2036 Gene Ontology (GO) Terms for all outliers identified in Los Angeles (LA). Outputs for each region are stored in the excel spreadsheet GOTerms_full.xlsx

**Supp Table 3:** 670 Gene Ontology (GO) Terms for all outliers identified in San Diego (SD). Outputs for each region are stored in the excel spreadsheet GOTerms_full.xlsx

**Supp Table 4:** 16 Gene Ontology (GO) Terms for all outliers identified in Victoria, B.C. (Vic). Outputs for each region are stored in the excel spreadsheet GOTerms_full.xlsx

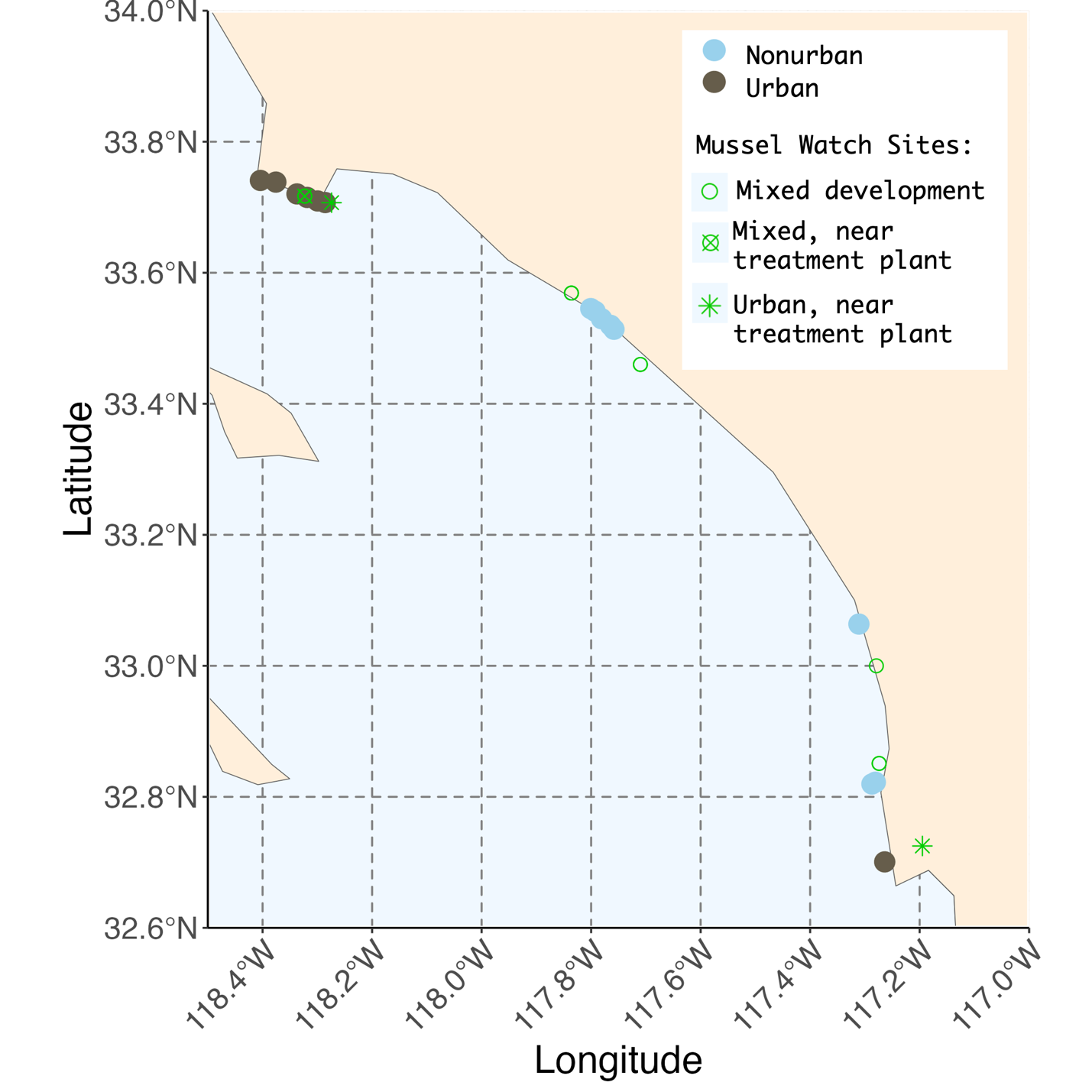

**Supp Fig 1**. Sampling map in relation to mussel watch collection sites. Colors correspond to classification of the Los Angeles (top) and San Diego (bottom) sites: nonurban (light blue) and urban (brown). Green shapes represent mussel watch site classifications. These sites were classified by land use, treatment plant proximity, and pollutant concentrations in *Mytilus* tissue and sediment (Maruya, Dodder, Schaffner, et al., 2014). We specifically highlight mixed development and urban sites in proximity to our sampling sites.

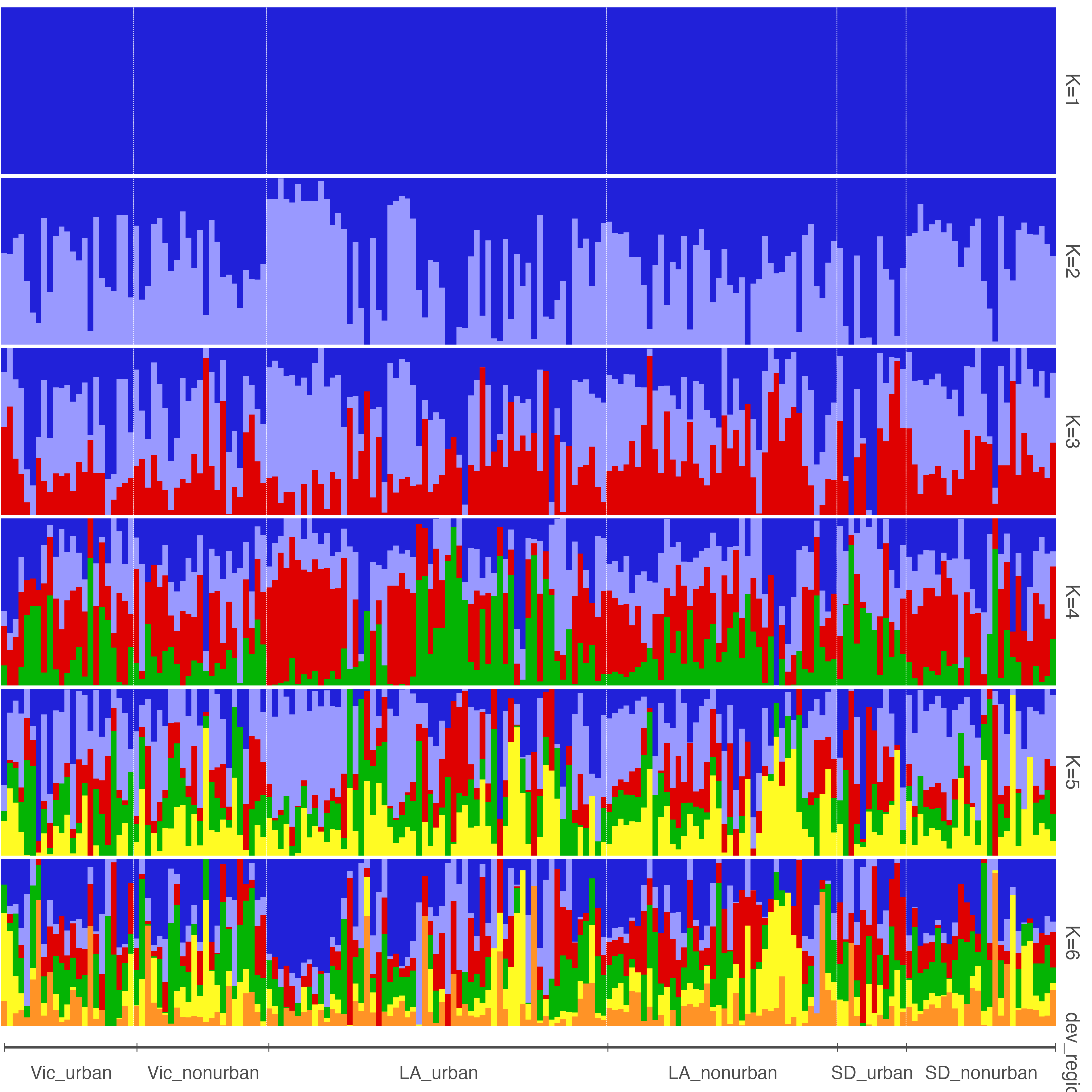

**Supp Fig 2**. Structure plots for K=1 through K=6 with hierarchical clustering implemented in Tess3r (v. 1.1.0) and Pophelper (v. 2.3.1) for visualization. The samples are organized by region (Victoria, Los Angeles and San Diego), and within each region the samples are separated by urban and nonurban sites. From top to bottom you can see increasing K values. The K that was the best fit for our data was K=1 (top panel).

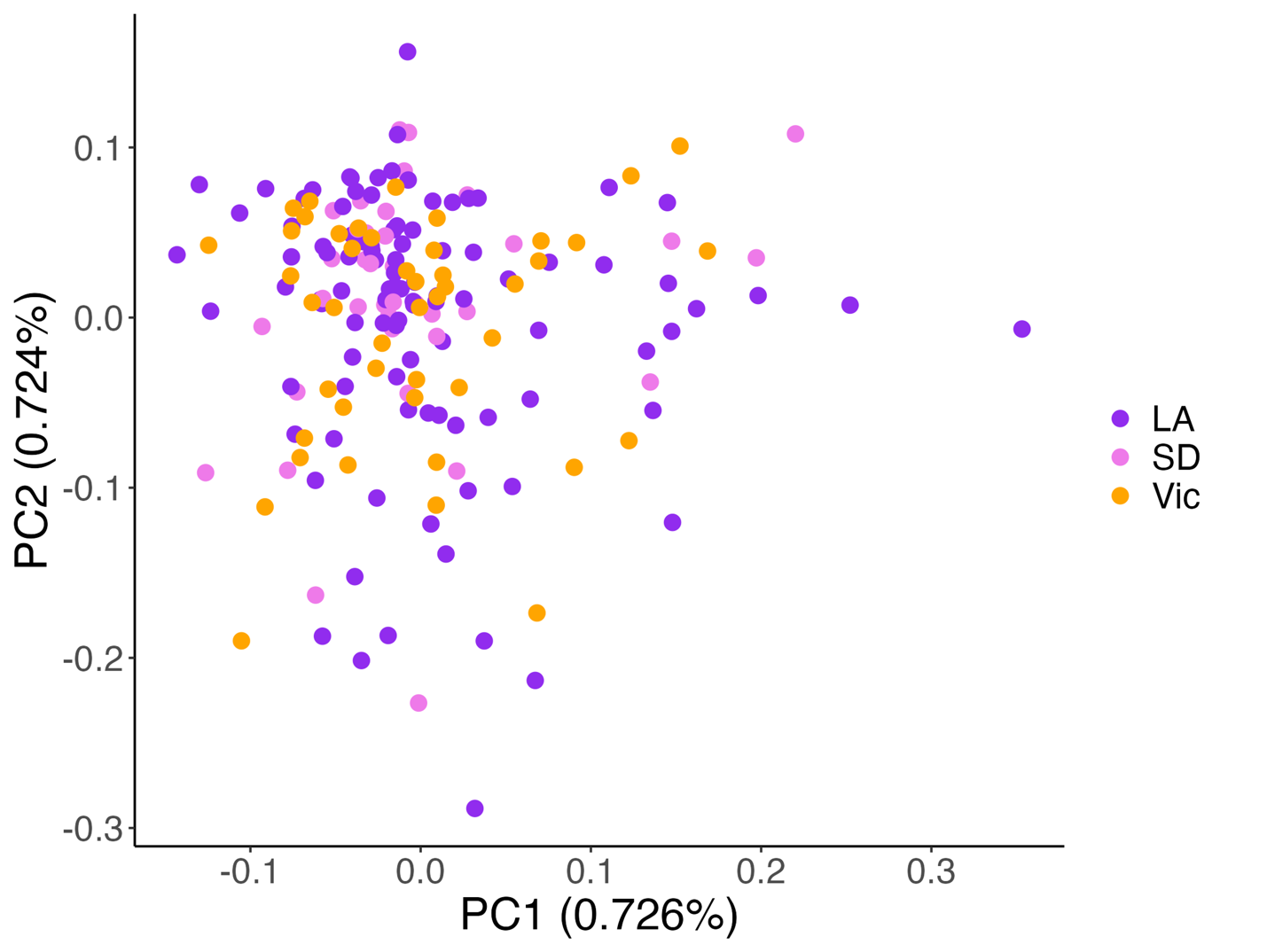

**Supp Fig 3.** We used a thinned set of 19,299 SNPs to analyze population structure, reducing effects of linkage disequilibrium. Thinned PCA colored by region: Los Angeles (LA) is purple, San Diego (SD) is pink and Victoria, B.C. (Vic) is orange. The axes correspond to principle components capturing 0.726% of the variation in the data (PC1) and 0.724% of the variation in the data (PC2). There is no clear grouping by region in this PCA.

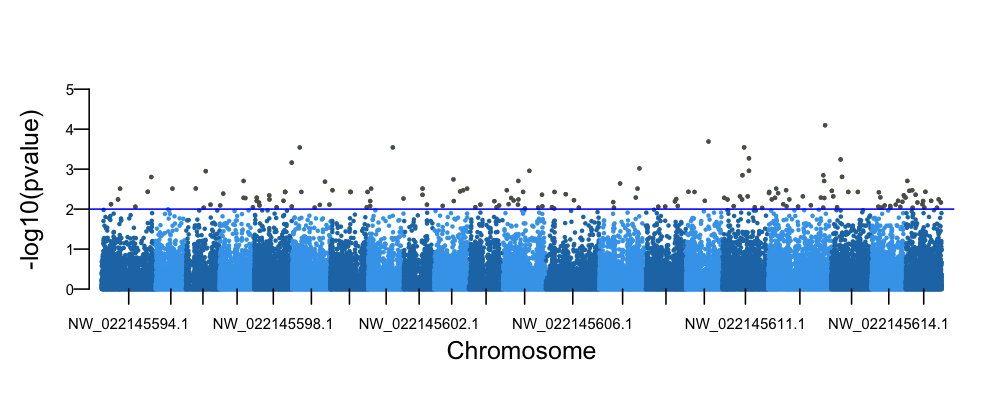

**Supp Fig 4.** Manhattan Plot using OutFLANK (v. 0.2) to identify outliers associated with urban status. The blue line indicates an outlier cutoff of p<0.01. There are 165 outliers associated with urban status, shown as grey dots above the blue line.

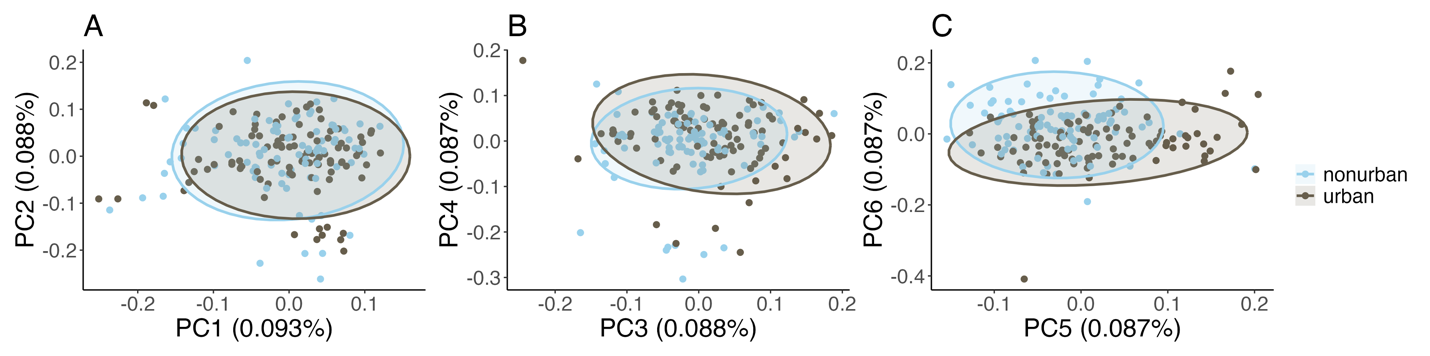

**Supp Fig 5.** Principle components (PCs) were identified using *pcadapt* (v. 4.4.0). PC1-PC6 shown here. Urban sites are shown in brown, while nonurban sites are shown in light blue. Urban and nonurban samples significantly differentiated along several PC axes: PC3 (E, t-value=2.941, p=0.0037), PC5 (F, t-value= 4.997, p<0.001), and PC6 (F, t-value= -4.586, p<0.001).

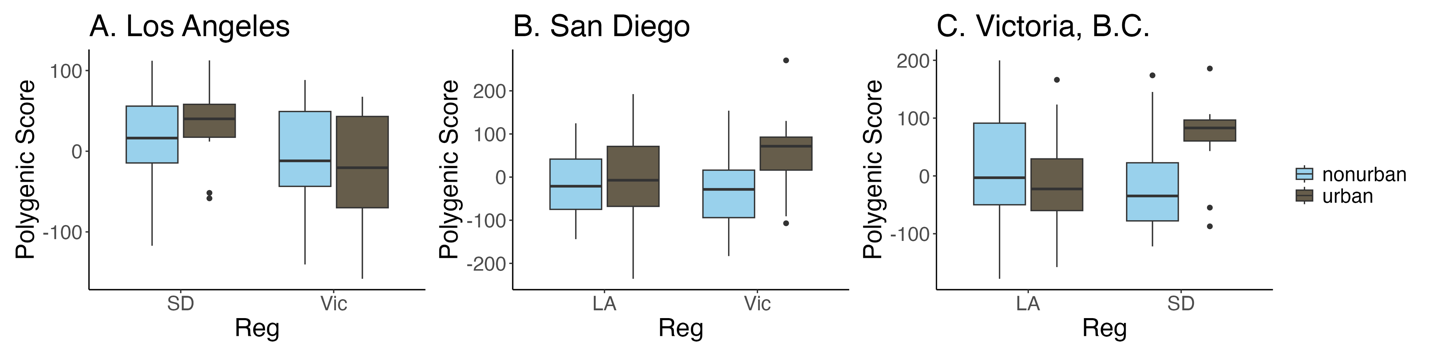

**Supp Fig 6.** To test the predictive power of one region versus the others for detecting differences between urban/nonurban samples, region and the interaction, we compared one region to the remaining two for all three comparisons. The training region is listed at the top of each boxplot for A-C and the compared regions are boxplots and labeled on the x-axis. The three regions are San Diego (SD), Victoria, B.C. (Vic) and Los Angeles (LA), and colors distinguish urban (brown) and nonurban (light blue) samples. (A) When comparing Los Angeles versus San Diego and Victoria, urban (two-way ANOVA, F_1,1_= 0.341, p=0.561) and the interaction of urban and region (two-way ANOVA, F_1,1_= 1.465, p=0.230) were not significant, while region was significant (two-way ANOVA, F_1,1_= 5.721, p=0.019). (B) When comparing San Diego versus Los Angeles and Victoria, urban (two-way ANOVA, F_1,1_= 6.845 p=0.010) and the interaction of urban and region (two-way ANOVA, F_1,1_= 5.493, p=0.020) were significant. However region was not (two-way ANOVA, F_1,1_= 1.976 p=0.162) (C) Lastly, when comparing Victoria versus Los Angeles and San Diego, neither region (two-way ANOVA, F_1,1_= 0.750, p=0.388) nor urban (two-way ANOVA, F_1,1_= 0.079, p=0.779) were significant, but the interaction of the two was (two-way ANOVA, F_1,1_= 9.699, p=0.002).

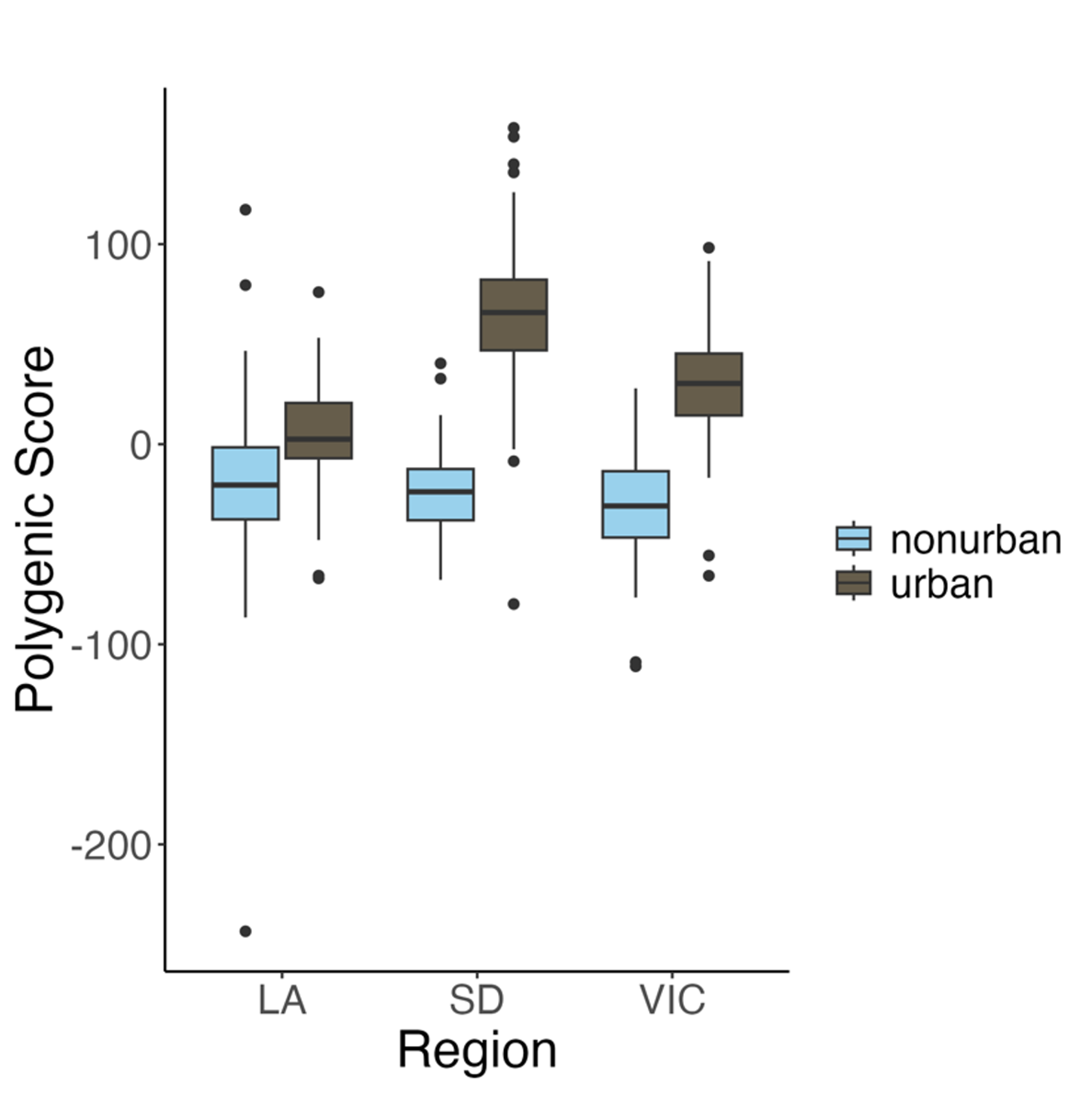

**Supp Fig 7.** To test the predictive power of our polygenic score test while accounting for sample size variation between regions, we conducted a downsampled PGS. The three regions are San Diego (SD), Victoria, B.C. (Vic) and Los Angeles (LA), and colors distinguish urban (brown) and nonurban (light blue) samples. 71/100 runs were significant for this analysis.
